# Protist and Fungal Metaxin-like Proteins: Relationship to Vertebrate Metaxins that Function in Protein Import into Mitochondria

**DOI:** 10.1101/2021.01.05.425400

**Authors:** Kenneth W. Adolph

## Abstract

Metaxin-like proteins are shown to be encoded in the genomes of a wide range of protists and fungi. The metaxin proteins were originally described in humans and mice, and were experimentally demonstrated to have a role in the import of nascent proteins into mitochondria. In this study, metaxin-like proteins of protists and fungi predicted from genome sequences were identified by criteria including their sequence homology with vertebrate metaxin proteins and the existence of distinctive metaxin protein domains. Protists of diverse taxa, including amoebae, protozoa, phytoplankton, downy mildews, water molds, and algae, were found to possess genes for metaxin-like proteins. With fungi, the important taxonomic divisions (phyla) of Ascomycota, Basidiomycota, Chytridiomycota, Mucoromycota, and Zoopagomycota had species with metaxin-like protein genes. The presence of distinctive GST_N_Metaxin, GST_C_Metaxin, and Tom37 domains in the predicted proteins indicates that the protist and fungal proteins are related to the vertebrate metaxins. However, the metaxin-like proteins are not direct homologs of vertebrate metaxins 1, 2, or 3, but have similarity to each of the three. The alignment of the metaxin-like proteins of a variety of protists with vertebrate metaxins 1 and 2 showed about 26% and 19% amino acid identities, respectively, while for fungal metaxin-like proteins, the identities were about 29% and 23%. The different percentages with the two vertebrate metaxins indicates that the metaxin-like proteins are both metaxin 1-like and, to a lesser degree, metaxin 2-like. The secondary structures of protist and fungal metaxin-like proteins both consist of nine α-helical segments, the same as for the vertebrate metaxins, with a negligible contribution from β-strand. Phylogenetic analysis demonstrated that the protist and fungal metaxin-like proteins and the vertebrate metaxins form distinct and separate groups, but that the groups are derived from a common ancestral protein sequence.

## 1. INTRODUCTION

Vertebrate metaxins 1 and 2 are proteins of the outer mitochondrial membrane with a role in the uptake of newly synthesized mitochondrial membrane proteins and their assimilation into the outer membrane. A metaxin 3 gene, originally detected in the zebrafish *Danio rerio* and the African clawed frog *Xenopus laevis*, has been demonstrated to be widely distributed among vertebrates (Adolph, 2019). Invertebrates have also been found to encode proteins related to vertebrate metaxins 1 and 2 (Adolph, 2020a). These include marine invertebrates such as the model research organisms *Ciona intestinalis* and *Aplysia californica*, and insects such as the fruit fly *Drosophila melanogaster* and the yellow fever mosquito *Aedes aegypti*. In addition, genes for metaxin-like proteins were revealed in plants such as *Arabidopsis thaliana*, a model plant in biological research, and bacteria such as *Pseudomonas aeruginosa*, responsible for hospital-acquired infections (Adolph, 2020b). Plant and bacterial genomes code for one, and sometimes more, metaxin-like proteins that are not directly homologous to vertebrate metaxins 1, 2, or 3. The purpose of the investigation reported here was to determine whether protists—eukaryotic organisms that are not animals, plants, or fungi—and a variety of fungi possess genes for metaxins or metaxin-like proteins. The results demonstrate that both protists and fungi encode genes for metaxin-like proteins. Important features of the predicted proteins are therefore described.

The protists with metaxin-like proteins that received the greatest emphasis in this study include model organisms and protists with fully sequenced genomes. Among these is *Chlamydomonas reinhardtii* (genome sequence: Merchant et al., 2007), a single-celled green alga used extensively in molecular biology and photosynthesis research. Another alga with a sequenced genome, the colonial green alga *Volvox carteri* (Prochnik et al., 2010), is a model for the evolution of multicellularity and cellular differentiation. An additional example is the phytoplankton *Emiliania huxleyi* (Read et al., 2013), responsible for large-scale marine algal blooms. The oomycete or water mold *Phytophthora infestans* (Haas et al., 2009) is a plant pathogen that caused the Irish Potato Famine of the 1840s. Two additional algae emphasized in this report are extremophiles: the red alga *Galdieria sulphuraria* (Schonknecht et al., 2013) is thermoacidophilic (grows well at high temperatures and low pH); *Dunaliella salina* (Polle et al., 2017) is halophilic (can survive in high salt concentrations) and has a red-orange color from possessing large amounts of β-carotene.

The investigation of fungi with metaxin-like proteins involved model organisms or organisms that are significant for human health or the environment, and with genomes that have been completely sequenced. An example is *Pneumocystis jirovecii* (genome sequence: Cisse et al., 2012), which causes *Pneumocystis* pneumonia in immunocompromised humans. *Penicillium chrysogenum,* another fungus with a sequenced genome (van den Berg et al., 2008), has been used in manufacturing the antibiotic penicillin. *Phycomyces blakesleeanus* (Corrochano et al., 2016), also in this report, is a model organism to investigate environmental signaling, particularly light sensing. The fungus *Aspergillus nidulans* (Galagan et al., 2005) is a well-established model organism for research in eukaryotic cell biology. Sequencing of fungal genomes has also been important for biochemical studies that include the methylation of fungal nucleic acids (Mondo et al., 2017). Single-cell genomics methods have enabled sequencing of the genomes of uncultured fungi such as the saprotroph *Blyttiomyces helicus* (Ahrendt et al., 2018), thus greatly expanding the information about uncultured fungi.

The metaxin 1 gene was initially identified as a mouse gene (*Mtx1*) that codes for a protein of the outer mitochondrial membrane. The protein was shown experimentally to function in the uptake of newly synthesized proteins into mitochondria (Armstrong et al., 1997). The human genome also possesses a metaxin 1 gene (*MTX1*), between the thrombospondin 3 gene (*THBS3*) and a glucocerebrosidase pseudogene (*psGBA1*) at cytogenetic location 1q21 (Long et al., 1996; Adolph et al., 1995). A metaxin 2 protein in the mouse was revealed by its binding to the metaxin 1 protein (Armstrong et al., 1999). Metaxin 2 might therefore interact with metaxin 1 in the cell, and function with metaxin 1 in the uptake of proteins into mitochondria. Mouse metaxin 1 and metaxin 2 proteins have substantial sequence differences, sharing only 29% amino acid identities. A homologous human gene (*MTX2*) at 2q31.2 is, like the mouse gene (*Mtx2*), next to a cluster of homeobox genes that code for transcription factors.

Metaxin 3, a metaxin distinct from metaxins 1 and 2, was initially revealed by the sequencing of cDNAs of the zebrafish *Danio rerio* (Adolph, 2004). The deduced metaxin 3 protein sequence of 313 amino acids is more homologous to zebrafish metaxin 1 than 2 (40% vs. 26% identical amino acids). Sequencing of cDNAs also led to the identification of a metaxin 3 gene in the African clawed frog *Xenopus laevis* (Adolph, 2005), like zebrafish a popular model vertebrate in biological research. As with zebrafish, the predicted *Xenopus* protein shows greater homology to metaxin 1 than to metaxin 2 (39% and 22%, respectively). Human metaxin 3 (Adolph, 2019) and zebrafish metaxin 3 proteins share 57% identical amino acids, while the human and *Xenopus* proteins share 67%, indicating the high degree of homology among these proteins.

The yeast system has provided the most detailed information about the uptake of proteins into mitochondria (Pfanner et al., 2019; Wiedemann and Pfanner, 2017; Neupert, 2015). Protein import into *Saccharomyces cerevisiae* mitochondria and incorporation into the outer mitochondrial membrane involves the TOM and SAM protein complexes. These protein complexes work together to bring about the uptake of nascent proteins, in particular β-barrel membrane proteins. Two conserved protein domains, the Tom37 domain and the Tom37_C domain, are major protein domains of the SAM complex. The vertebrate metaxins, and many metaxin-like proteins, also possess a Tom37 domain in addition to GST_N_Metaxin and GST_C_Metaxin domains (Adolph, this study). This is compatible with the suggestion that the metaxins function in mitochondrial protein import.

Previous studies (Adolph, 2020a,b) have indicated that the metaxin proteins are not confined to vertebrates, but metaxins and metaxin-like proteins are encoded in the genomes of a wide variety of organisms. Invertebrates, plants, and bacteria all possess genes for metaxins or metaxin-like proteins. Two other fundamental groups of organisms—protists and fungi—have therefore been investigated. The results demonstrate that the genomes of many protists and fungi code for metaxin-like proteins. Basic features of the deduced proteins are described and compared to the metaxins and metaxin-like proteins of other organisms.

## 2. METHODS

Identifying the conserved protein domains of the metaxin-like proteins of protists and fungi (Figure 1) used the CD_Search tool (www.ncbi.nlm.nih.gov/Structure/cdd/wrpsb.cgi) of the NCBI (National Center for Biotechnology Information)(Lu et al., 2020). Alignment of protein sequences as in Figure 2 to determine amino acid identities and similarities employed the NCBI Global Alignment tool to compare the full lengths of two sequences (https://blast.ncbi.nlm.nih.gov/Blast.cgi; Needleman and Wunsch, 1970; Altschul et al., 1990). Also used was Align Two Sequences BLAST to align the protein regions of highest homology. In addition, pairwise sequence alignments were carried out with the EMBOSS Needle tool (https://www.ebi.ac.uk/Tools/psa/emboss_needle/). The α-helical secondary structures of metaxin-like proteins were predicted using the PSIPRED server (bioinf.cs.ucl.ac.uk/psipred/; Jones, 1999). To compare the α-helical segments of selected proteins (Figure 3), their amino acid sequences were aligned with the NCBI COBALT multiple sequence alignment tool (www.ncbi.nlm.nih.gov/tools/cobalt/; Papadopoulos and Agarwala, 2007). Evolutionary relationships among metaxin-like proteins were also investigated with the COBALT tool; phylogenetic trees, as shown in Figure 4, were generated from the alignments. Also used was Clustal Omega (https://www.ebi.ac.uk/Tools/msa/clustalo/; Sievers et al., 2011). Transmembrane α-helical segments were detected using the PHOBIUS server (https://phobius.sbc.su.se; Kall et al., 2007), and, in addition, the TMHMM server (www.cbs.dtu.dk/services/TMHMM/; Krogh et al., 2001). Identification of genes near metaxin-like genes was provided by “Gene neighbors” information from protein BLAST searches. Adjacent genes were also identified with the Genome Data Viewer and Map Viewer genome browsers (www.ncbi.nlm.nih.gov/genome/gdv/).

**Figure 1.**
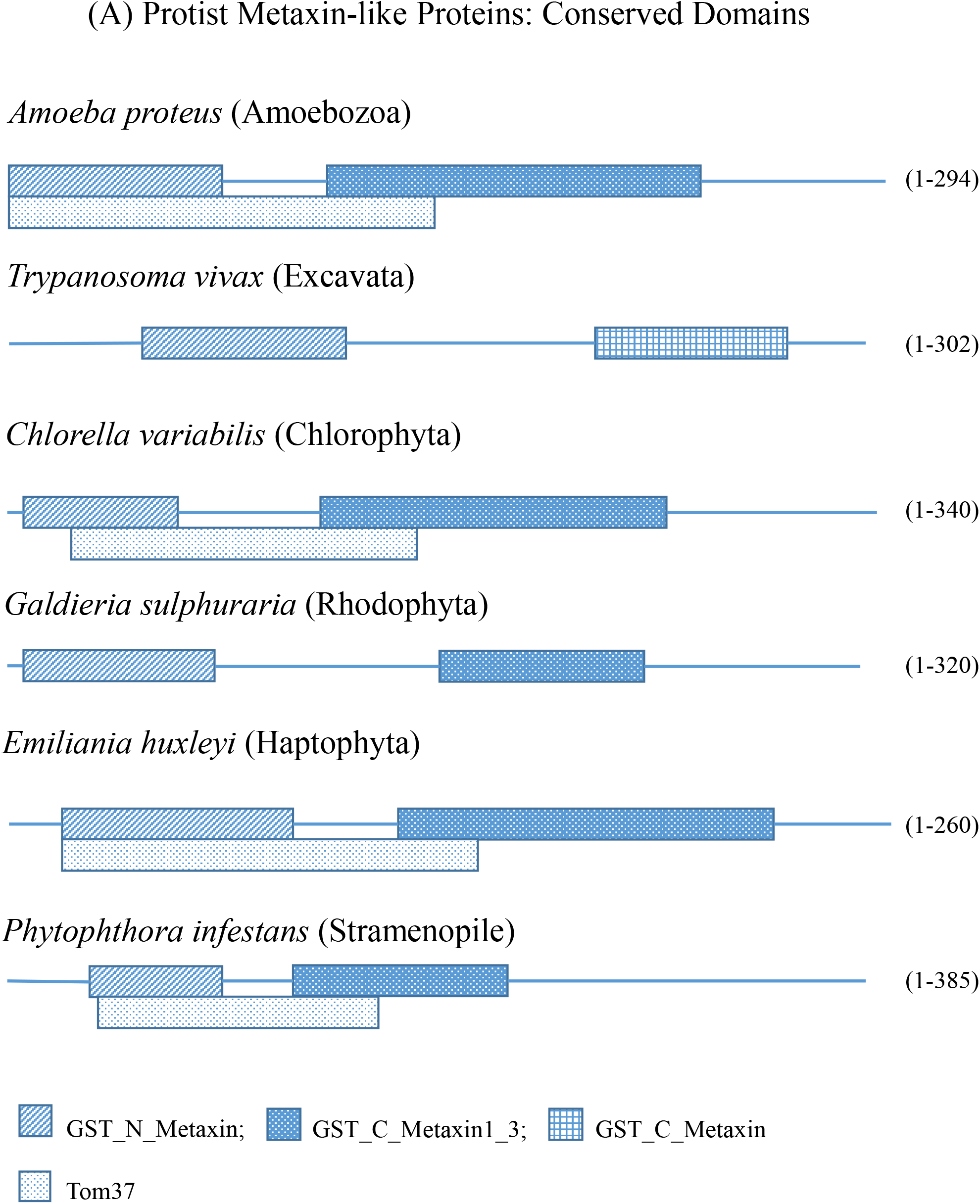
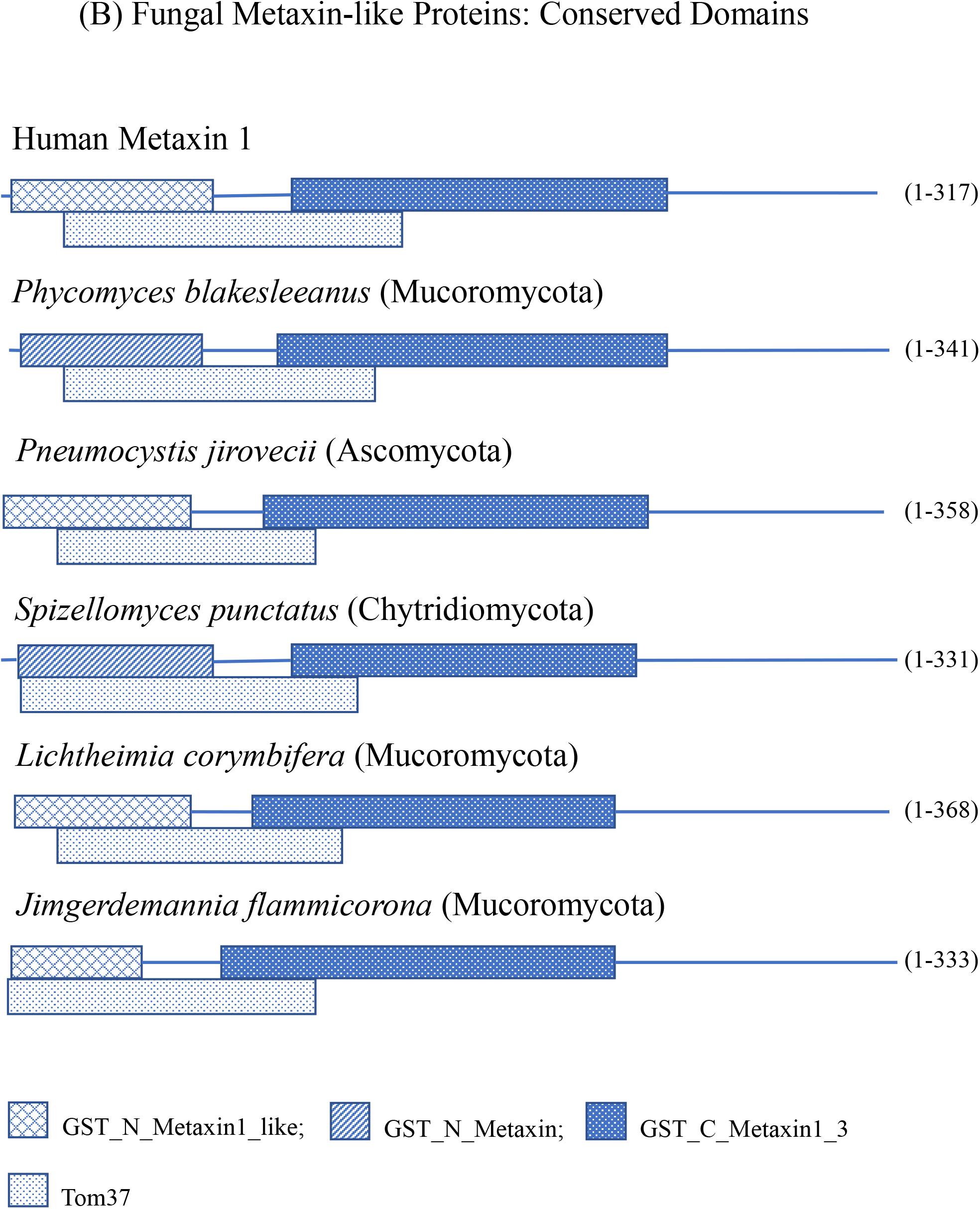
Domain structures of protist and fungal metaxin-like proteins. (A) The figure shows the characteristic GST_N_Metaxin and GST_C_Metaxin domains of the metaxin-like proteins of representative protists. The Tom37 domain, originally described in yeast mitochondrial import proteins, is also included. The GST_Metaxin domains are distinguishing features of vertebrate and invertebrate metaxins, and plant and bacterial metaxin-like proteins. The protists in Figure 1A include *Amoeba proteus*, an amoeba of the phylum Amoebozoa, *Trypanosoma vivax*, a protozoan of the supergroup Excavata, and *Chlorella variabilis*, a green alga of the phylum Chlorophyta. Also in the figure are *Galdieria sulphuraria*, a red alga of the phylum Rhodophyta. *Emiliania huxleyi*, a phytoplankton of the phylum Haptophyta, and *Phytophthora infestans*, a water mold of the Stramenopile lineage. (B) The GST_N_Metaxin, GST_C_Metaxin, and Tom 37 domains of the metaxin-like proteins of select fungi are included in the figure. The conserved protein domains of human metaxin 1 are also shown for comparison with the fungal proteins. The representative fungal metaxin-like proteins are those of *Phycomyces blakesleeanus*, a model fungus of the taxonomic division or phylum Mucoromycota, *Pneumocystis jirovecii* (previously *carinii*), a human pathogen of the Ascomycota, and *Spizellomyces punctatus*, a soil fungus of the Chytridiomycota. Also in the figure are two additional members of the Mucoromycota, *Lichtheimia corymbifera*, an opportunistic human pathogen, and *Jimgerdemannia flammicorona*, a fungal symbiont of plant roots.

**Figure 2.**
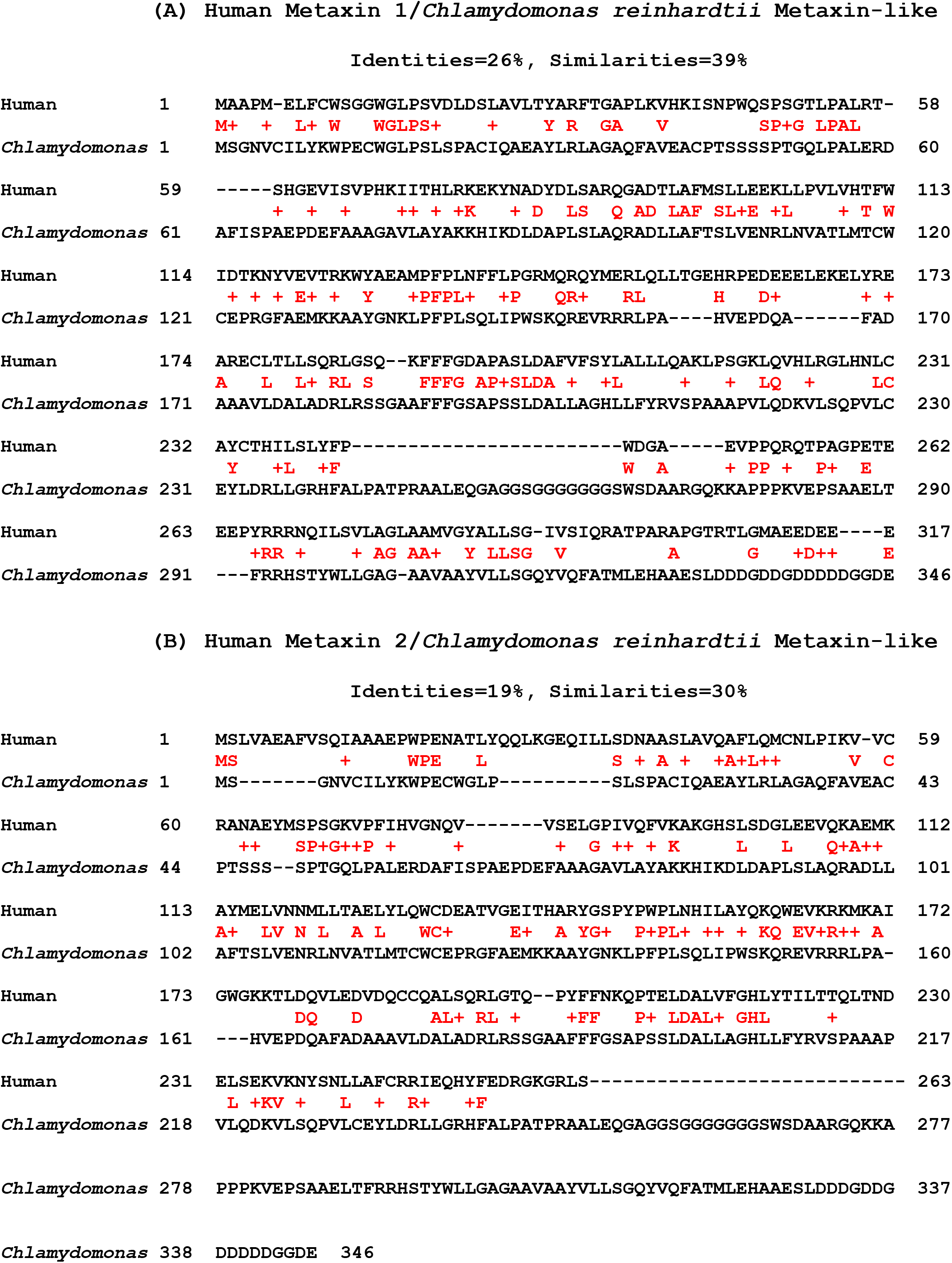
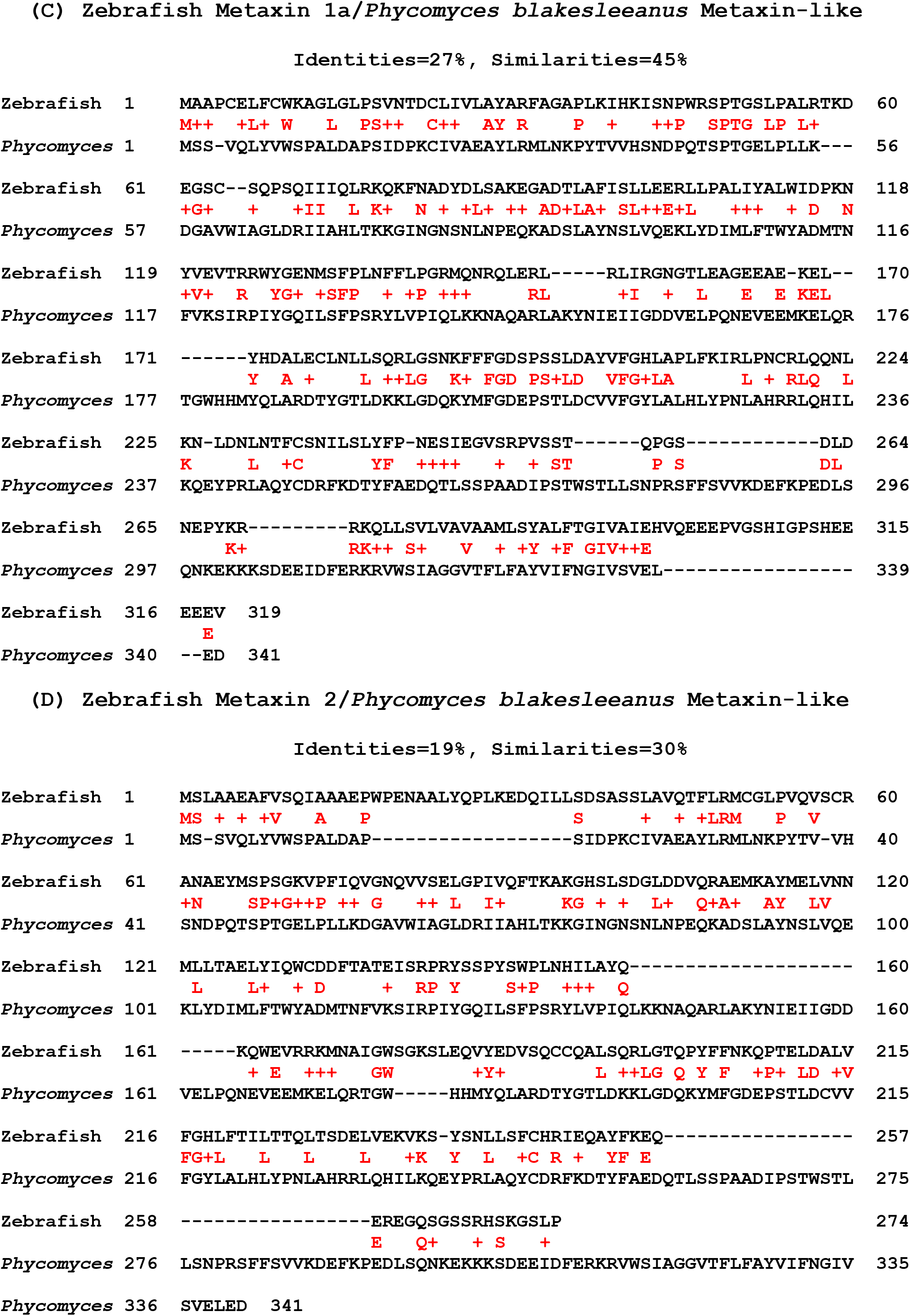
Identities and similarities of the amino acid sequences of protist and fungal metaxin-like proteins. (A, B) The amino acid sequence of the metaxin-like protein of *Chlamydomonas reinhardtii* is aligned with the sequences of human metaxin 1 (Figure 2A) and human metaxin 2 (Figure 2B). *Chlamydomonas* is a widely used model organism in cell and molecular biology research, including photosynthesis research. Between the aligned sequences are the identical amino acids, shown as single-letter abbreviations in red, and similar amino acids, shown by + signs. For human metaxin 1, the identical amino acids are 26% and the similar amino acids are 39% (Figure 2A). For human metaxin 2, the identities and similarities are 19% and 30% (Figure 2B). The *Chlamydomonas* protein has 346 amino acids, whereas human metaxin 1 has 317 and human metaxin 2 has 263. These percentages were obtained by comparing the entire lengths of the sequences using BLAST Global Align. The percentages using BLAST Align Two Sequences, which only include regions of high homology, are about 6-8% higher. (C, D) The sequence of the *Phycomyces blakesleeanus* metaxin-like protein is aligned with the protein sequences of zebrafish metaxin 1a and zebrafish metaxin 2. *Phycomyces* is a model fungus for investigating the responses of fungi to environmental stimuli. In the figure, the identical and similar amino acids are between the aligned protein sequences, highlighted in red. Aligned with zebrafish metaxin 1a, the *Phycomyces* metaxin-like protein has 27% identical amino acids (Figure 2C); zebrafish metaxin 2 and the fungal protein share 19% identical amino acids (Figure 2D). The *Phycomyces* protein has 341 residues, while zebrafish metaxin 1a has 319 and zebrafish metaxin 2 has 274. The alignments with the protist (*Chlamydomonas*) and fungal (*Phycomyces*) metaxin-like proteins in Figure 2 demonstrate that the greatest homology is to vertebrate metaxin 1, but with significant homology to vertebrate metaxin 2.

**Figure 3.**
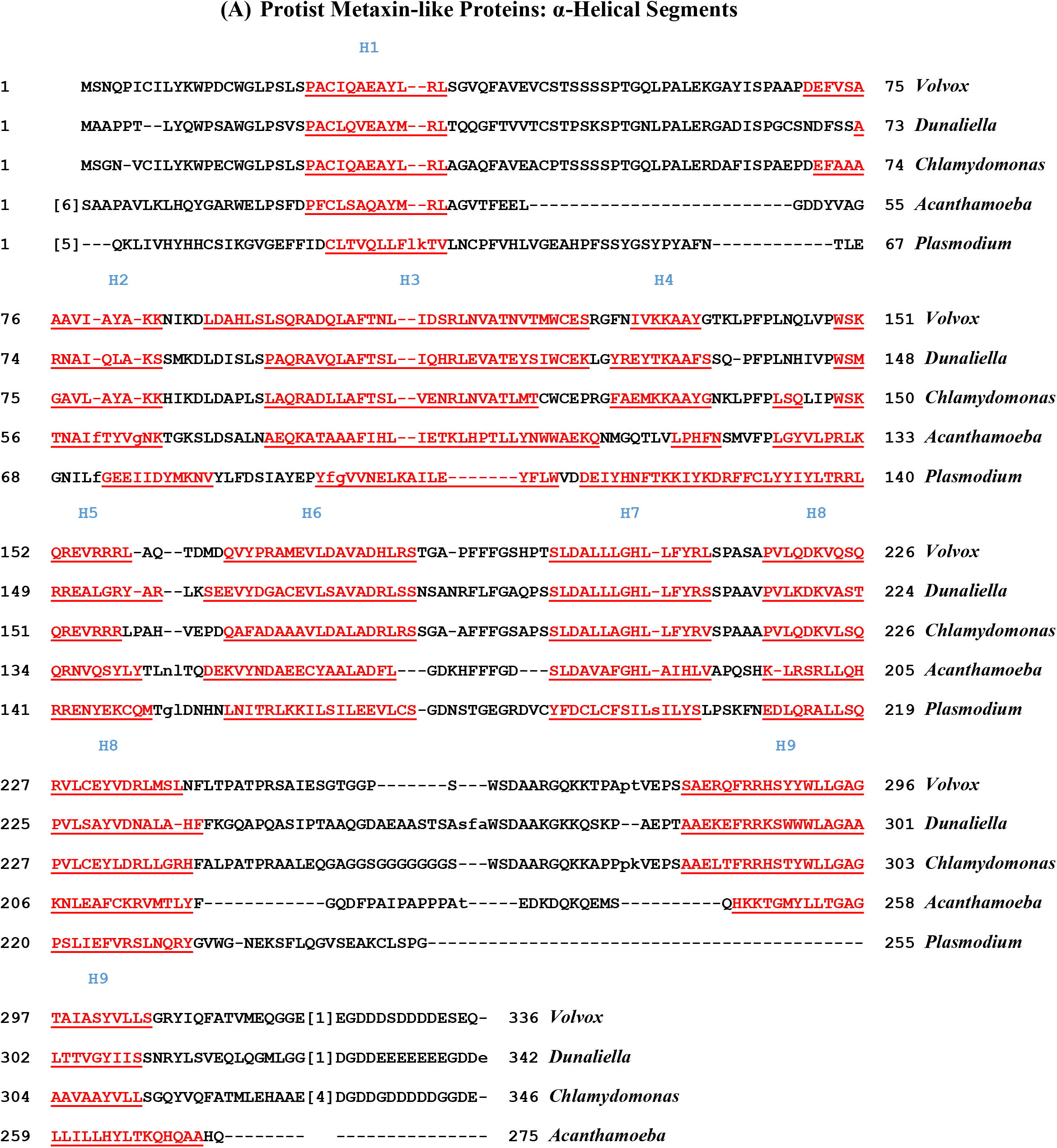
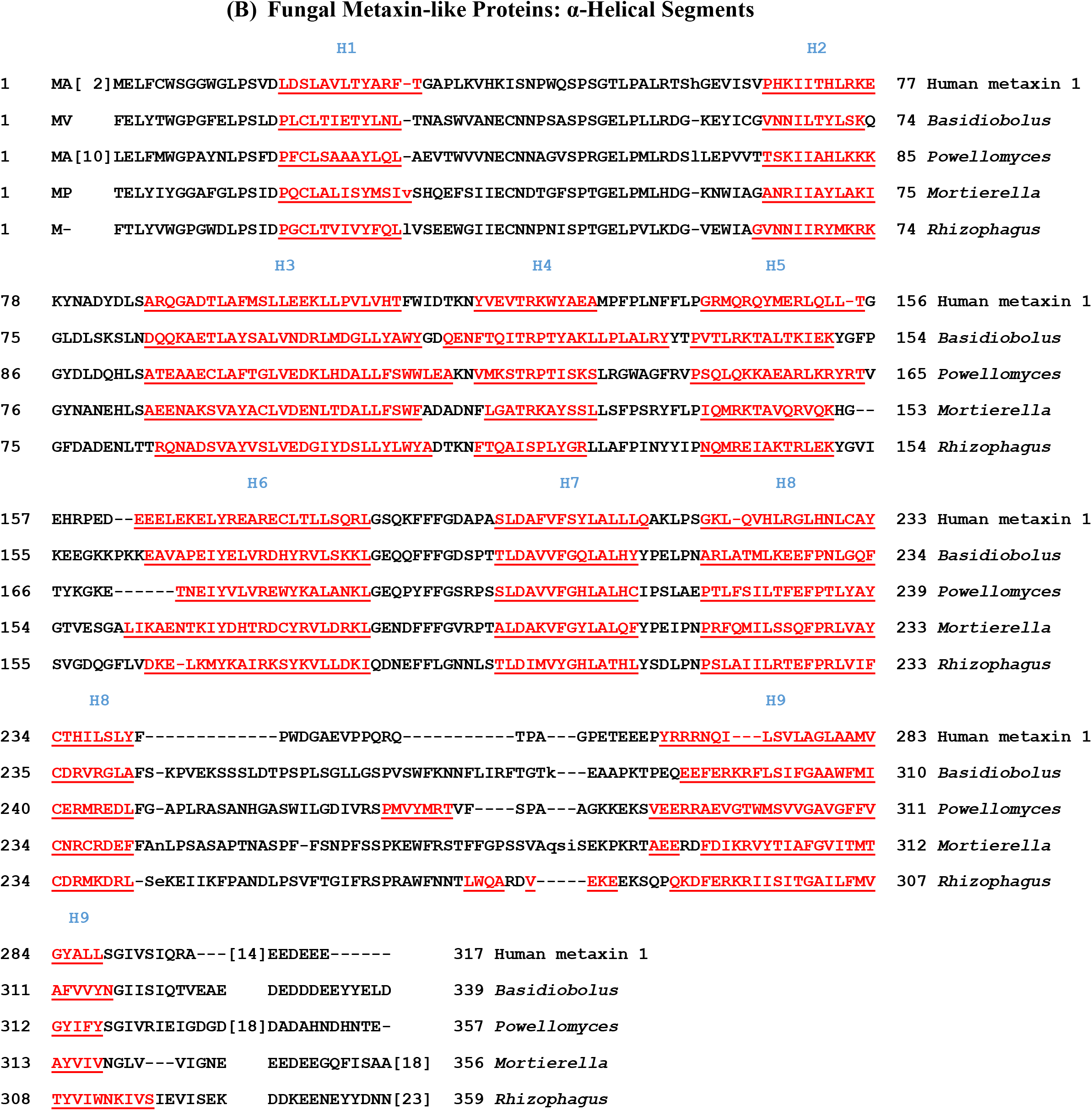
Alpha-helical segments of metaxin-like proteins of protists and fungi: multiple sequence alignments. (A) The predicted α-helical segments of the metaxin-like proteins of five selected protists are shown in red and underlined. The secondary structures of the proteins consist primarily of α-helical segments, labeled H1 to H9. The figure includes three chlorophyte algae, *Volvox carteri*, *Dunaliella salina*, and *Chlamydomonas reinhardtii*, as well as the amoebozoan *Acanthamoeba castellanii* and the apicomplexan parasite *Plasmodium vivax*. The number of α-helices and their locations along the amino acid chains are highly conserved between the different protist species. The same pattern was also found, with small variations, for many other protist metaxin-like proteins that were investigated. In addition, the number of helices and spacings between helices also characterize vertebrate metaxin proteins, in particular human metaxins 1 and 3. Human metaxin 2 is almost identical, but lacks helix H9 and has an extra N-terminal helix. In Figure 3B, which shows metaxin-like proteins of fungi, the number of α-helical segments and the spacings between segments are strikingly similar to the results in (A) for representative protists. The four fungal metaxin-like proteins and human metaxin 1 in (B) all have nine α-helical segments, H1 through H9, as was found for the protists. This observation helps to confirm the identification of the fungal proteins as metaxin-like, and indicates that metaxin-like proteins are a widely distributed group of proteins across a spectrum of living organisms. The fungi included in the figure are *Basidiobolus meristosporus* (Division: Zoopagomycota or Entomophthoromycota), *Powellomyces hirtus* (Chytridiomycota), *Mortierella verticillata* (Mucoromycota), and *Rhizophagus irregularis* (Mucoromycota or Glomeromycota).

**Figure 4.**
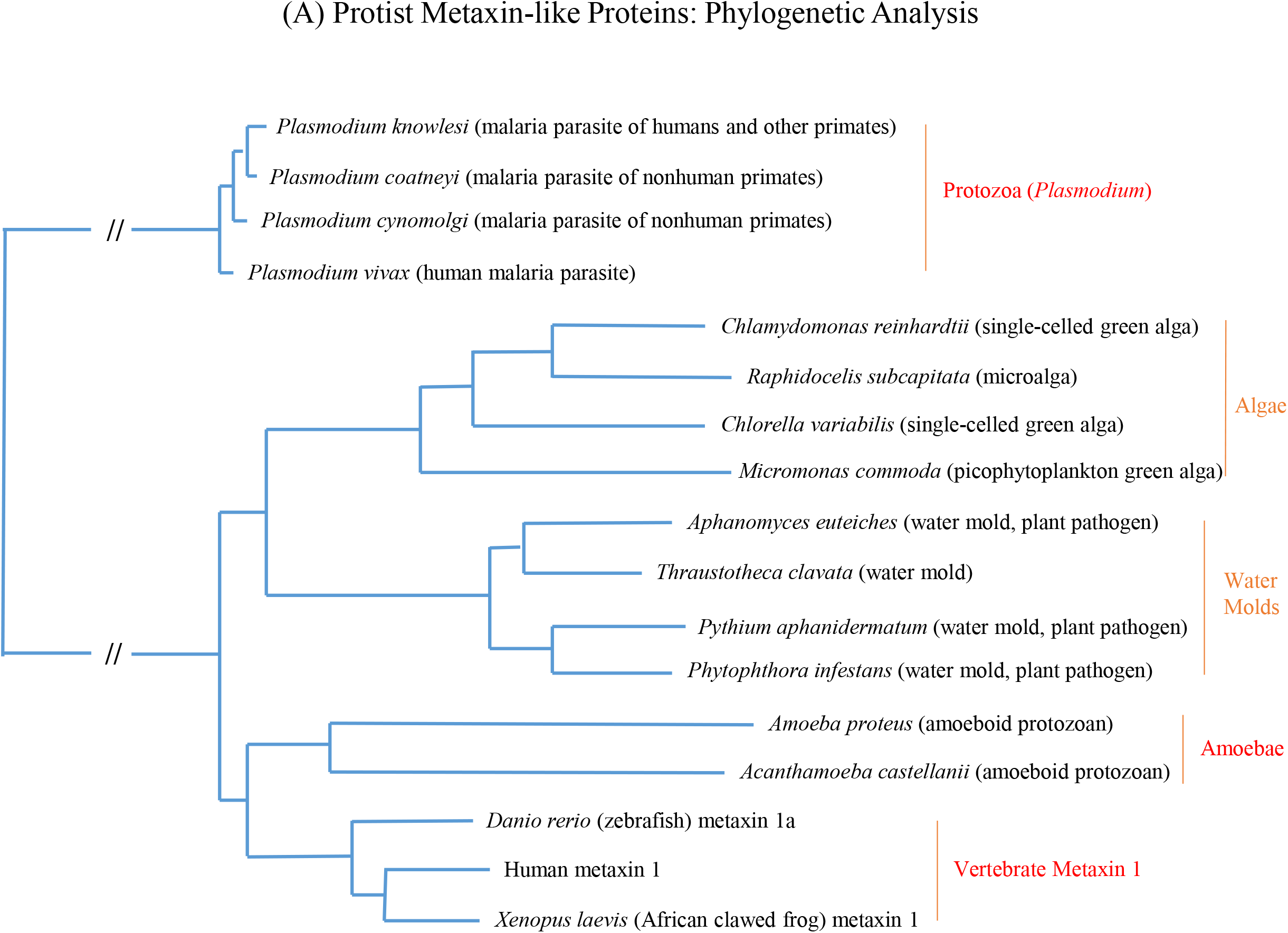
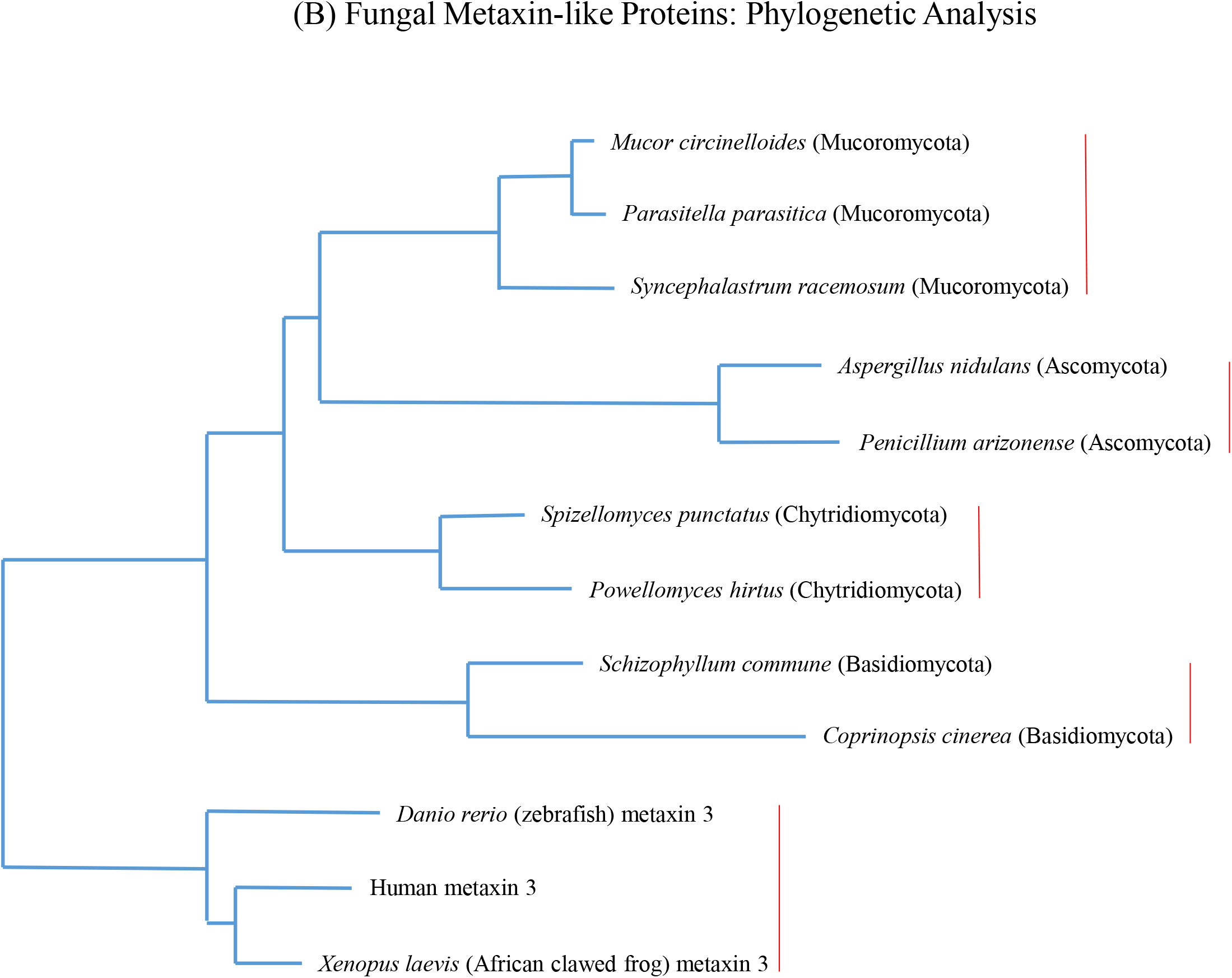
Phylogenetic analysis of protist and fungal metaxin-like proteins. (A) The figure shows the evolutionary relationships of the metaxin-like proteins of a variety of protists. These include algae, water molds, amoebae, and protozoa. The horizontal lengths of the branches indicate the extent of evolutionary change. The metaxin-like proteins of different protist taxa form distinct clusters on the phylogenetic tree. The figure demonstrates that the metaxin-like proteins of algae, water molds, and amoebae are related to vertebrate metaxins, in particular vertebrate metaxin 1. The vertebrate metaxin 1 proteins—human, *Xenopus*, and zebrafish—form a separate cluster that is phylogenetically connected to the clusters of the protist metaxin-like proteins. A more distant relationship is found for the *Plasmodium* metaxin-like proteins, which only possess GST_C_Metaxin domains, and not GST_N_Metaxin domains. (B) The phylogenetic tree for representative fungal metaxin-like proteins displays the clustering of the fungi of the same taxonomic division (phylum). The *Mucor*, *Parasitella*, and *Syncephalastrum* fungi are all in the Mucoromycota division, and form a cluster due to being related by evolution. The same is true of the Ascomycota fungi, *Aspergillus* and *Penicillium*, the Chytridiomycota fungi, *Spizellomyces* and *Powellomyces*, and the Basidiomycota fungi, *Schizophyllum* and *Coprinopsis*. In each case, the metaxin-like proteins of fungi of the same division are grouped together, indicating they are related by evolution. The vertebrate metaxin 3 proteins in the figure also cluster together, and, most importantly, are seen to be related to the fungal metaxin-like proteins.

## 3. RESULTS AND DISCUSSION

### 3.1. Presence of Metaxin-like Proteins in Protists and Fungi

To investigate whether protist and fungal genomes encode metaxin-like proteins, protein sequence databases of the NCBI and JGI (DOE Joint Genome Institute) were searched with vertebrate metaxin protein sequences. Predicted proteins homologous to vertebrate metaxins were revealed for a variety of protist and fungal species. The identity of the proteins as metaxin-like proteins was strengthened by the existence in the proteins of GST_N_Metaxin and GST_C_Metaxin protein domains. These domains are distinguishing features of vertebrate metaxins, and also invertebrate metaxins (Adolph, 2020a) and plant and bacterial metaxin-like proteins (Adolph, 2020b). Other properties of the metaxin-like proteins, such as the lengths of the protein chains, had to be consistent with the properties of vertebrate and invertebrate metaxins.

For the protists, a single metaxin-like protein was encoded, similar to the situation in plants and bacteria, instead of distinct metaxin 1, metaxin 2, and metaxin 3 proteins as in vertebrates. Metaxin-like proteins were found to be most common in algae, particularly green algae of the phylum Chlorophyta. These include *Chlamydomonas reinhardtii*, a model organism of cell and molecular biology research. Other well-studied chlorophyte algae found to have metaxin-like proteins are species of *Chlorella* and *Volvox*. Metaxin-like proteins were identified as well in red algae of the phylum Rhodophyta and in algae of the phylum Cryptophyta and the phylum Charophyta. Phytoplankton and the fungus-like water molds and downy mildews are additional protists with species that have genes for metaxin-like proteins. Protists that are animal-like such as amoebae and other protozoa were also found to encode metaxin-like proteins.

However, metaxin-like proteins were not detected for a number of types of protists. These include choanoflagellates, ciliates like *Paramecium* and *Tetrahymena*, diatoms, dinoflagellates, foraminifera, *Giardia*, and slime molds like *Dictyostelium discoideum* and *Physarum polycephalum*. Metaxin-like proteins may be found for these protists as more complete genome sequences become available.

The existence of metaxin-like proteins in fungi was revealed, as for the protists, through database searches with the protein sequences of vertebrate metaxins, in particular the human metaxins. Predicted fungal metaxin-like proteins that correspond to vertebrate metaxins 1, 2, and 3 were not found. Instead, the fungal proteins have significant homology to all three vertebrate metaxins, with the greatest homology to metaxin 1. The fungal divisions (phyla) with species that were found to possess metaxin-like proteins include the Ascomycota, Basidiomycota, Chytridiomycota, Mucoromycota, and Zoopagomycota. Ascomycota fungi that encode metaxin-like proteins include the human pathogen *Pneumocystis jirovecii* (previously *carinii*). A Basidiomycota fungus with a metaxin-like protein is the mushroom *Schizophyllum commune*, and a Chytridiomycota fungus with the protein is the plant pathogen *Synchytrium microbalum*. The Mucoromycota division, that has perhaps the largest number of fungi possessing genes for metaxin-like proteins, is represented by the model fungus *Phycomyces blakesleeanus*, and the Zoopagomycota division by the human pathogen *Basidiobolus meristosporus* (*ranarum*).

### 3.2. Conserved Protein Domains of Protist and Fungal Metaxin-like Proteins

The possession of GST_N_Metaxin and GST_C_Metaxin protein domains is a feature that defines the metaxin proteins and metaxin-like proteins. In addition, the Tom37 domain is a common feature of vertebrate metaxins including metaxin 3 (Adolph, 2019), invertebrate metaxins 1 and 2 (Adolph, 2020a), and bacterial metaxin-like proteins (Adolph, 2020b). In this study, protists of various taxa were found to contain a single metaxin-like protein, like plants and bacteria, and not homologs of the vertebrate and invertebrate metaxins. Figure 1A shows the protein domains of the metaxin-like proteins of a selection of protists of different phyla. The protists include *Amoeba proteus*, a large amoeboid protozoan, *Trypanosoma vivax*, a parasitic protozoan, *Chlorella variabilis*, an algal model for carbon assimilation in plants, *Galdieria sulfuraria*, a red alga tolerant of high temperatures and low pH, *Emiliania huxleyi*, an abundant phytoplankton, and *Phytophthora infestans*, a plant pathogen. The conserved domains provide strong evidence that these proteins, and homologous proteins of protists not in the figure, are metaxin-like proteins.

Fungal metaxin-like proteins, included in Figure 1B along with human metaxin 1, are also characterized by GST_N_Metaxin, GST_C_Metaxin, and Tom 37 conserved protein domains. No other conserved domains are significant features of the fungal proteins. The lengths of the protein chains, such as the examples in Figure 1B, are generally similar to the vertebrate metaxins, e.g. 317 and 326 amino acids for human metaxins 1 and 3, respectively. However, some fungal metaxin-like proteins, typically with an extended C-terminal sequence, can exceed 400 amino acids. Examples are *Aspergillus nidulans* with 421 residues and *Penicillium arizonense* with 435 residues. The fungal metaxin-like proteins in Figure 1B are those of *Phycomyces blakesleeanus*, studied for its strong phototropism response, *Pneumocysti*s *jirovecii* (*carinii*), a human pathogen that causes *Pneumocystis* pneumonia, *Spizellomyces punctatus*, a soil fungus significant in terrestrial ecosystems, *Lichtheimia corymbifera*, an opportunistic pathogen in humans with impaired immunity, and *Jimgerdemannia flammicorona*, a fungus used in research into the symbiotic relationships of fungi and plants. The conserved protein domains of fungi resemble those of protists (Figure 1A), vertebrate metaxins such as human metaxin 1 (Figure 1B), and metaxin-like proteins (Adolph, 2020a,b).

### 3.3. Homology of Protist and Fungal Metaxin-like Proteins: Alignment of Amino Acid Sequences

Amino acid alignments demonstrate that the metaxin-like proteins of protists are more homologous to human metaxin 1 than to human metaxin 2. But the difference is not major, and the protist metaxin-like proteins have substantial homology to human metaxin 2. Alignments with human metaxin 3 also show substantial homology, similar to human metaxin 1. The example in Figure 2 shows that the metaxin-like protein of *Chlamydomonas reinhardtii*, an important model organism in biology research, has 26% identical amino acids aligned with human metaxin 1 (Figure 2A) and 19% with human metaxin 2 (Figure 2B). The percentages of similar amino acids are 39% and 30%, respectively. Comparable results are found in aligning the *Chlamydomonas* protein with other vertebrate metaxins. In particular, the *Chlamydomonas* metaxin-like protein has 28% identities aligned with zebrafish (*Danio rerio*) metaxin 1a and 20% aligned with zebrafish metaxin 2. For *Xenopus laevis* metaxin 1, 27% identities are found, while for *Xenopus* metaxin 2, 18%. The results with *Chlamydomonas* are also true for other protists, including amoebae, phytoplankton, downy mildews, water molds, and algae. As examples, for the *Amoeba proteus* metaxin-like protein, the identities are 24% with human metaxin 1 and 20% with human metaxin 2, and for the phytoplankton *Emiliania huxleyi*, the percentages are 28% and 21%.

The amino acid identities and similarities in comparing one protist metaxin-like protein with another depend upon the taxonomic relatedness of the protists. Consider the *Chlamydomonas reinhardtii* metaxin-like protein sequence aligned with the sequences of other green algae of the same phylum, Chlorophyta. With *Gonium pectorale* 68% identities are found, with *Monoraphidium neglectum* 43%, and with *Chlorella variabilis* 40%. When metaxin-like proteins of different protist phyla are aligned, the amino acid identities are significantly lower and average about 22%.

For the fungi, Figure 2 shows the alignment of the metaxin-like protein of the fungus *Phycomyces blakesleeanus* with zebrafish metaxin 1a (Figure 2C) and metaxin 2 (Figure 2D). *Phycomyces* is widely-used in biological research into responses to environmental stimuli. The most studied of these stimuli is light, but other stimuli include gravity, physical contact, air movement (wind), and nearby objects (avoidance response). With zebrafish metaxin 1a, the identical amino acids are 27%, and with zebrafish metaxin 2, 19%. Aligning other vertebrate metaxins with the *Phycomyces* metaxin–like protein strengthens the observation that the metaxin-like protein is most homologous to vertebrate metaxin 1, but with significant homology to metaxin 2. As an example, human metaxin 1 aligned with the *Phycomyces* protein reveals 29% identities, while human metaxin 2 reveals 25% identities. With *Xenopus laevis* metaxin 1 and metaxin 2, the identities are 28% and 26%, respectively. Similar results are found when other metaxin-like proteins of fungi of different taxonomic divisions are aligned with vertebrate metaxins. For instance, the metaxin-like protein of the human pathogen *Pseudomonas jirovecii* (Division: Ascomycota) shows 30% identities aligned with zebrafish metaxin 1a, the soil fungus *Spizellomyces punctatus* (Chytridiomycota) also shows 30% identities, as does the human pathogen *Basidiobolus meristosporus* (Zoopagomycota). In each of these three cases, the identities with metaxin 2 are lower.

The percentages of identical amino acids are substantially higher when comparing the *Phycomyces* metaxin-like protein sequence with those of other fungi of the same taxonomic division, Mucoromycota. As examples, the *Phycomyces* protein shows 69% identities when aligned with the protein of the thermophilic fungus and opportunistic human pathogen *Lichtheimia corymbifera*, and 63% identities when aligned with the protein of the soil fungus and emerging human pathogen *Mucor circinelloides*.

Table 1 lists some basic features of the predicted amino acid chains of representative metaxin-like proteins. These include numbers of amino acids, molecular weights, and percentages of acidic, basic, and hydrophobic residues. For the protists, the different taxonomic groups include Charophyta, Chlorophyta, Cryptophyta, and Stramenopiles. The numbers of amino acids for the protist metaxin-like proteins are generally similar to those for human metaxins 1 and 3 (317 and 326, respectively). Some have longer chains, such as the *Saprolegnia* protein, due to having a longer C-terminal region that extends beyond the GST_C_Metaxin domain. Metaxin-like proteins of fungi are also included in Table 1. These examples represent major taxonomic divisions (phyla) of the fungi: Ascomycota, Basidiomycota, Chytridiomycota, Mucoromycota, and Zoopagomycota. Like the protist proteins, the lengths of the amino acid chains of the fungal proteins are similar to those of the vertebrate metaxins.

**Table 1.**
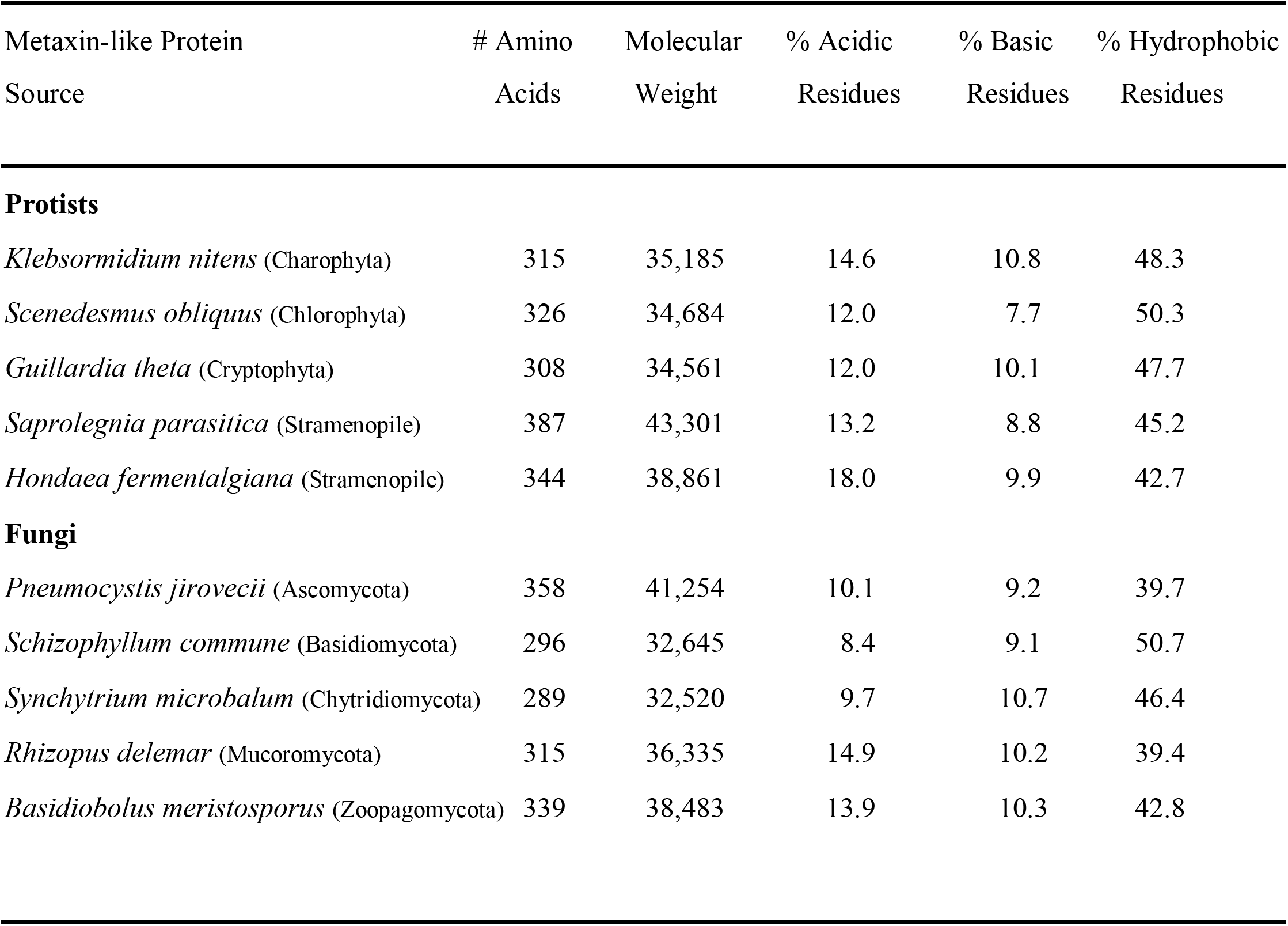
Amino Acid Analysis of the Metaxin-like Proteins of Protists and Fungi

### 3.4. Alpha-Helical Secondary Structures of Protist and Fungal Metaxin-like Proteins

The predicted secondary structures of the metaxin-like proteins of selected protists are shown in Figure 3A. The protists in the multiple-sequence alignment include *Volvox carteri*, a colonial (multicellular) green alga, *Dunaliella salina*, a halophilic (high salt tolerant) green alga, *Chlamydomonas reinhardtii*, a model green alga of biology research, *Acanthamoeba castellanii*, also a model organism of biology research, and *Plasmodium vivax*, a human malaria parasite. Segments of α-helix, underlined and colored red, predominate in the patterns of secondary structure for the five examples, with very little β-strand. The same pattern of α-helical segments was found for a large number of other protists of diverse types—amoebae, protozoa, phytoplankton, downy mildews, water molds, algae. The patterns of α-helix are made up of nine segments, H1 to H9, with similar spacings between pairs of helices. The number of helical segments and the spacings between segments are identical to the patterns found for metaxins 1 and 3 of humans and other vertebrates (Adolph, 2019). Vertebrate metaxin 2 has eight segments, H1 to H8, with an extra ninth segment near the N-terminal end of the protein.

The vertebrate metaxin 1 and 2 patterns of helices are also found for invertebrate metaxins 1 and 2 (Adolph, 2020a). Plant metaxin-like proteins have the vertebrate metaxin 1 pattern of nine helices (Adolph, 2020b). Bacterial metaxin-like proteins have eight helices (Adolph, 2020b), without the extra N-terminal helix of metaxin 2. The conservation of helices is especially striking since many of the vertebrate and invertebrate metaxins, and plant and bacterial metaxin-like proteins, share no more than about 20% amino acid identities. It demonstrates that the α-helical structures of metaxins and metaxin-like proteins are more highly conserved than their amino acid sequences.

A characteristic feature of vertebrate metaxin 1 is a predicted transmembrane α-helical segment near the C-terminal end. Many protist metaxin-like proteins were also found in this study to have a transmembrane helix close to the C-terminus. The transmembrane helix largely overlaps the final α-helical segment, H9, in Figure 3. *Chlamydomonas reinhardtii*, for example, has a C-terminal transmembrane helix between amino acid residues 295 and 312 of the 346 amino acid protein. *Amoeba proteus*, another example, has a predicted transmembrane helix between amino acids 269 and 289 of the 294 amino acid protein. The existence of a C-terminal transmembrane helix was most commonly observed for algae. But not all protist metaxin-like proteins were found to have C-terminal transmembrane helices. The significance of the presence or absence of the transmembrane helix may become clearer as additional metaxin-like protein sequences of protists become available for analysis.

Figure 3B includes multiple-sequence alignments of the metaxin-like proteins of four representative fungi, as well as human metaxin 1. Shown in red and underlined are the predicted α-helical segments. The fungal species are *Basidiobolus meristosporus*, the cause of subcutaneous infections (zygomycosis) in humans, *Powellomyces hirtus*, a soil fungus, *Mortierella verticillata*, a fungus that grows on decaying organic matter (saprotroph), and *Rhizophagus irregularis*, a fungus that forms a symbiotic association with plant roots (mycorrhiza). The secondary structures of the fungal proteins, like the protist proteins in Figure 3A, consist of nine α-helices, labeled H1 through H9. β-strand is mainly not present. The spacings between the α-helices are also similar to the protist metaxin-like proteins. This is remarkable given that alignments of fungal with protist metaxin-like proteins generally show less than 25% amino acid identities.

Metaxin-like proteins of fungi of different taxonomic divisions—Ascomycota, Basidiomycota, Chytridiomycota, Mucoromycota, Zoopagomycota—were also found to possess a transmembrane helix near the C-terminus. The helix largely coincides with the final α-helical segment, H9. For instance, the metaxin-like protein of *Pneumocystis murina* (Ascomycota), a pathogenic fungus of mice, has a transmembrane helix between residues 315-335 of the 359 amino acid chain. *Rhizopus delemar* (Mucoromycota), a food mold and potential human pathogen, is another fungus that has a metaxin-like protein with a transmembrane helix (residues 272-295 of the 315 residue protein).

In vertebrates, the C-terminal transmembrane helix of metaxin 1 was hypothesized to anchor the protein to the outer mitochondrial membrane. The existence of C-terminal transmembrane helices for the protist and fungal metaxin-like proteins may also be connected with their roles as proteins of the outer mitochondrial membrane.

### 3.5. Evolutionary Relationships of Protist and Fungal Metaxin-like Proteins

Figure 4A shows the evolutionary relationships of representative protist metaxin-like proteins. The phylogenetic analysis demonstrates that the metaxin-like proteins of different types of protists—algae, water molds, amoebae, protozoa—form separate groups based on their taxonomic similarity. The groups are related by evolution from a common ancestor. The protists in the figure include green algae such as *Chlamydomonas reinhardtii*. Prominent among the water molds in the phylogenetic tree is *Phytophthora infestans*, the causative agent of the potato blight responsible for famines in the 1800s. Amoebae are represented by *Amoeba proteus* and *Acanthamoeba castellanii*. In addition, the figure includes *Plasmodium vivax*, which causes the most common form of malaria, and three other *Plasmodium* parasites.

Within each group in Figure 4A, protist metaxin-like proteins with a similar taxonomic classification are seen to be the most closely related. To illustrate, all four algae belong to the same phylum, Chlorophyta. *Chlamydomonas* and *Raphidocelis*, the most closely related of the algal metaxin-like proteins on the phylogenetic tree, also belong to the same taxonomic class, Chlorophyceae. *Chlorella* is in the class Trebouxiophyceae, also a core Chlorophyta class along with Chlorophyceae. *Micromonas* is in the more distantly related class Mamiellophyceae. The water molds in Figure 4A provide another example. The *Aphanomyces* and *Thraustotheca* water molds are Stramenopiles, Order Saprolegniales. And, correspondingly, the phylogenetic tree shows that they are the most closely related of the four water molds. The *Pythium* and *Phytophthora* water molds are also Stramenopiles, but are in a different order, Peronosporales.

Figure 4A also includes vertebrate metaxin 1 proteins (human, zebrafish, and *Xenopus*), which form a separate cluster from the protist metaxin-like proteins. The vertebrate metaxins are seen to be related by evolution to the protist proteins. The metaxins and metaxin-like proteins are all likely to have genes derived from a common ancestral gene. The *Plasmodium* metaxin-like proteins in the figure show a lower degree of relatedness to the vertebrate metaxins and other metaxin-like proteins. This is consistent with the *Plasmodium* proteins possessing only the GST_C_Metaxin protein domain, and lacking the GST_N_Metaxin domain characteristic of the metaxins.

The phylogenetic tree of fungal metaxin-like proteins in Figure 4B demonstrates that the fungal proteins of the same taxonomic division (phylum) are the most closely related. The figure includes metaxin-like proteins of four fungal divisions—Ascomycota, Basidiomycota, Chytridiomycota, and Mucoromycota. The three Mucoromycota fungi show the connection between taxonomic and evolutionary similarity. The *Mucor* and *Parasitella* proteins are in the same taxonomic order, Mucorales, and the same family, Mucoraceae, besides the same division, Mucoromycota. The *Syncephalastrum* protein, less related to the other two, has the same Mucorales order, but a different family, Syncephalastraceae. Vertebrate metaxin 3 proteins are also part of the phylogenetic tree, with the human and *Xenopus* proteins being more closely related than zebrafish metaxin 3.

### 3.6. Genes Adjacent to the Metaxin-like Genes of Protists and Fungi

The neighboring genes of protist metaxin-like proteins are generally not conserved between different protists, based on the limited annotation of protist genomes currently available. And none of the genomic regions next to the metaxin-like genes are homologous to the genomic regions of vertebrate metaxins. To give examples of the neighboring genes in protists, for *Acanthamoeba castellanii* the gene order is adenosylmethionine decarboxylase/GATA zinc finger domain containing protein/**metaxin-like protein**/tRNA intron endonuclease. For the green alga *Coccomyxa subellipsoidea*, the adjacent genes are different: phosphoenolpyruvate carboxylase/WD40 repeat-like protein/**metaxin-like protein**/glyceraldehyde-3-phosphate dehydrogenase/phosphoglycerate kinase. Exceptions to having different neighboring genes were found for genes of metaxin-like proteins that are very similar taxonomically. As more annotated protist genomes become available, further information should emerge about the significance of the differences in neighboring genes.

With fungal metaxin-like proteins, some similarities but mainly differences in neighboring genes are observed. For instance, the neighboring genes of the *Aspergillus fumigatus* (Ascomycota) metaxin-like protein are: zinc ion transporter/**metaxin-like protein**/allergen/ carboxyphosphoenolpyruvate phosphomutase. This partly resembles the gene order for *Penicillium digitatum* (Ascomycota): zinc ion transporter/**metaxin-like protein**/ allergen/oxaloacetate hydrolase. But for *Rhizophagus irregularis* (Mucoromycota), the neighboring genes are very different: **metaxin-like protein**/endonuclease V/tRNA-dihydrouridine synthase/cytochrome c oxidase. Relatively few fungal genomes have been well-annotated, and this limits the conclusions that can be made about neighboring genes.

## Notes

### Competing Interest Statement

The authors have declared no competing interest.

